# Protruding Geometry Provides Potential Homing Sites of Circulating Tumor Cells

**DOI:** 10.64898/2026.10.16.759535

**Authors:** Yeqing Leng, Bo Li

## Abstract

Circulating tumor cells (CTCs) must adhere to the vascular endothelium to initiate metastasis, yet the role of vessel wall topography remains unclear. We present an integrated microfluidic platform combining defined protruding geometries, live-cell imaging, and controlled perfusion to study CTC homing. Using a multiscale device that reproduces hierarchical vascular branching, we show that CTCs accumulate preferentially at locations with protruded geometry, including bifurcations, sharp corners and positively curved segments. At these hotspots, a viscoelastic, polymer-like network assembles, capturing CTCs and accompanying progressive multicellular aggregation. This network undergoes stretching and elastic recoil even under laminar flow. Mechanistically, knockdown of DSG2 and ITGB3 reduces retention, suggesting a potential anti-metastasis strategy targeting molecular–mechanical coupling. Computational fluid dynamics simulations indicate that hotspots coincide with pronounced velocity and pressure perturbations. Our experiments identify local geometry as a critical determinant of CTC arrest and provide a platform for studying the physical mechanisms of metastatic homing.

## Introduction

Most cancer-related deaths arise from metastatic dissemination, and circulating tumor cells (CTCs) constitute the blood-borne population that drives this process^1^. For a CTC to seed a distant organ, it must first contact and adhere to the vascular endothelium at a site permissive for stable arrest—the initial phase of homing^2^, a process that proceeds largely through mechanical trapping in vivo^3^. Although the biochemical regulators of CTC–endothelium interaction, including selectins, integrins, and chemokines, have been extensively characterized^4,5^, the physical topography of the vessel wall has received less attention^6^. Protrusions within the vascular lining—stenotic ridges, irregular endothelial contours, or engineered microstructures—can reconfigure local flow, producing stagnation zones, recirculation eddies, and steep wall-shear-stress gradients^7,8^. Such flow patterns may prolong cell–wall contact and favor adhesive bond formation, suggesting that geometric irregularities can serve as local homing sites^9^.

Direct evidence that protruding geometry drives CTC accumulation and the underlying biochemical-mechanical coupling nonetheless remains scarce. A principal limitation is the lack of experimental systems capable of simultaneously maintaining tumor cell viability, applying well-defined flow, and resolving adhesion events in real time with spatial precision^6^. Static adhesion and transwell formats cannot reproduce the dynamic shear environment of the vasculature, whereas endpoint microfluidic studies frequently overlook the transient contacts that precede stable arrest^8,10^.

To overcome these constraints, we devised an integrated platform (**Figure 1**). A microfluidic chip with defined protruding elements introduces controlled geometric disturbances into the flow channel. A fluorescence microscope (**Figure 1A**) is coupled to a live-cell platform containing this chip (**Figure 1B**), and the assembly is housed in an incubator with a fluidic unit, CO_2_ supply, and syringe pump, permitting prolonged culture and stable perfusion^11,12^. Tumor cells accumulate in discrete regions, and the spatial relationship between flow direction and the hotspot can be visualized directly (**Figure 1C**). Whereas previous studies described CTC dynamics in the circulation^8^ or arrest in a single fixed geometry^9^, this platform allows these variables to be manipulated independently, identifying geometric hotspots as in vitro proxies for homing sites^9^ and dissecting the coupling between flow and adhesion, extending beyond computational predictions of preferential adhesion at bifurcations and curved segments^13,14^. This work may illuminate a physical dimension of CTC arrest that has received little attention^8^ and could suggest new avenues for interfering with metastatic seeding.

**Figure 1.**
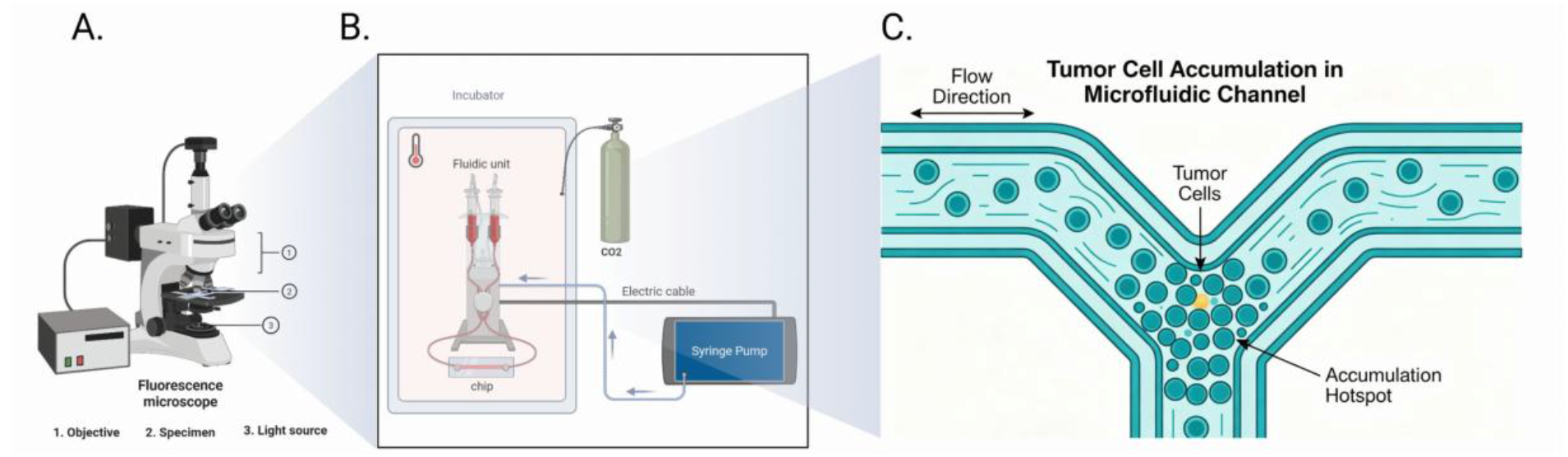
Schematic illustration of the experimental platform. A. Fluorescence microscope (1, objective; 2, specimen; 3, light source). B. Live-cell platform coupled to a microfluidic chip containing defined protruding elements; the assembly is housed in an incubator with a fluidic unit, CO_2_ supply, and syringe pump. C. Tumor cell accumulation in the microfluidic channel, with arrows indicating flow direction and the hotspot formed in relation to the protruding geometry. Created with BioRender.com.

Using this platform, we show that CTCs accumulate preferentially at protruding geometric confinements—bifurcations, sharp corners, and curved segments—within a multiscale vascular network (**Figure 2**). At these hotspots, a viscoelastic, polymer-like network assembles and captures CTCs through flow-induced stretching and elastic recoil, driving progressive accumulation (**Figure 3**). Molecular dissection reveals that DSG2 and ITGB3 mediate cell–cell and cell–matrix adhesion, while flow promotes multicellular aggregation (**Figure 4**). Computational fluid dynamics simulations further confirm that these hotspots coincide with pronounced velocity and pressure perturbations (**Figure 5**). Together, these findings identify local geometry as a critical determinant of CTC arrest and establish a platform for dissecting the physical mechanisms of metastatic homing.

**Figure 2.**
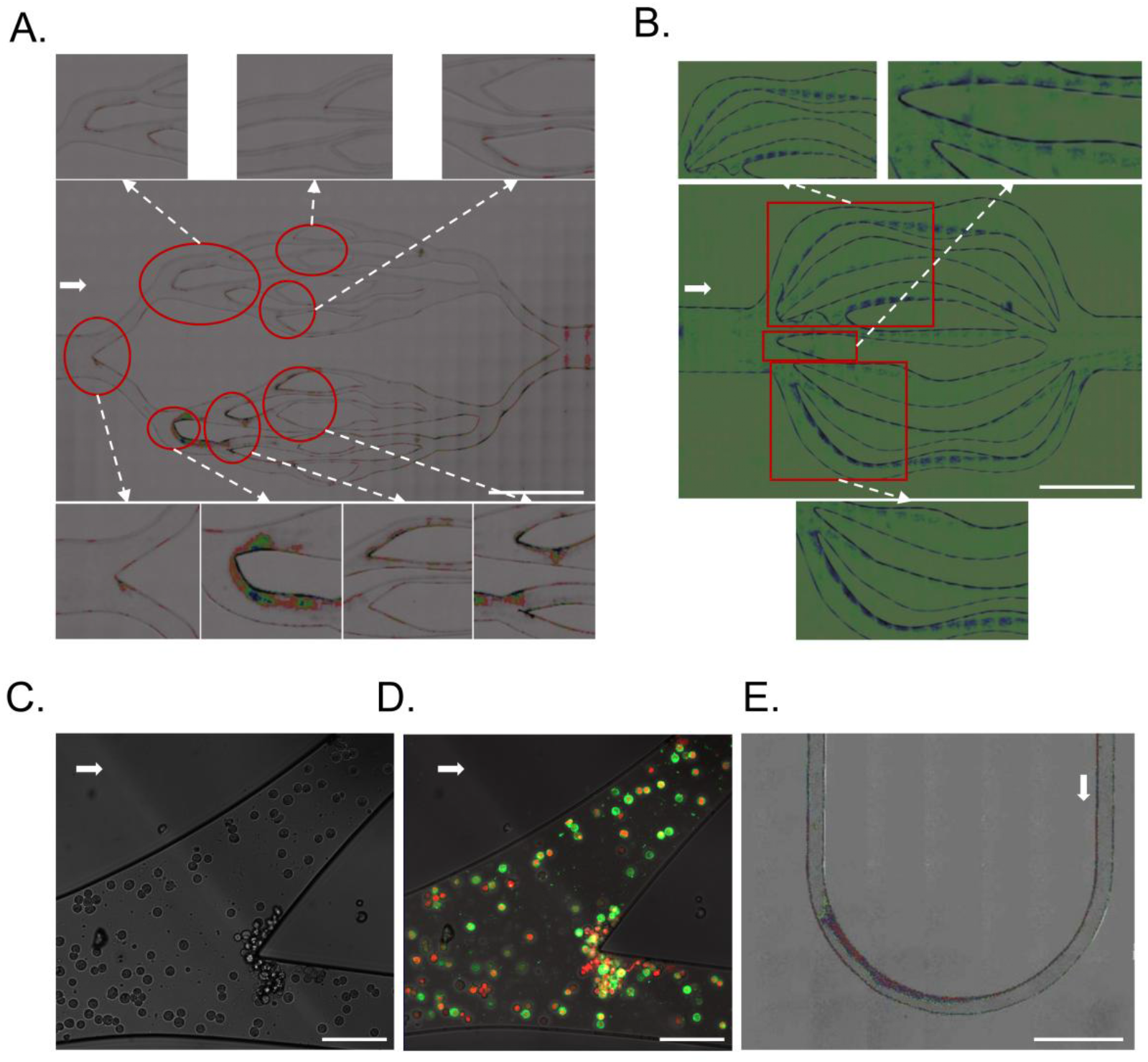
Flow-dependent spatiotemporal dynamics of tumor cells in hierarchically branched vascular networks. A. Microscopic images of the multiscale microfluidic chip, showing hierarchically branched channels (main vessel → secondary branches → capillary-like constrictions). Tumor cells are perfused under controlled flow. Scale bar: 2655 μm. B. Overview of the single-level vessel model showing tumor cell distribution within the main channel. Insets highlight representative hotspots of cell accumulation, adhesion, and aggregation. Scale bars: 2655 μm (overview) and as indicated (insets). C. Bright-field micrographs at branching points, showing tumor cell accumulation, adhesion, and flow recirculation zones at bifurcations, corners, and stenosis-like constrictions. Scale bar: 24.5 μm. D. Confocal micrographs at branching points, further demonstrating tumor cell accumulation and adhesion at bifurcations, corners, and stenosis-like constrictions, consistent with flow-induced hotspot formation. Scale bar: 24.5 μm. E. Bright-field micrograph of a curved channel segment within the microfluidic device, showing tumor cell accumulation at the bend. Cells preferentially accumulate and adhere along the inner wall of the curvature, indicative of flow-induced hotspot formation. Arrow indicates flow direction. Scale bar: 1100 μm.

**Figure 3.**
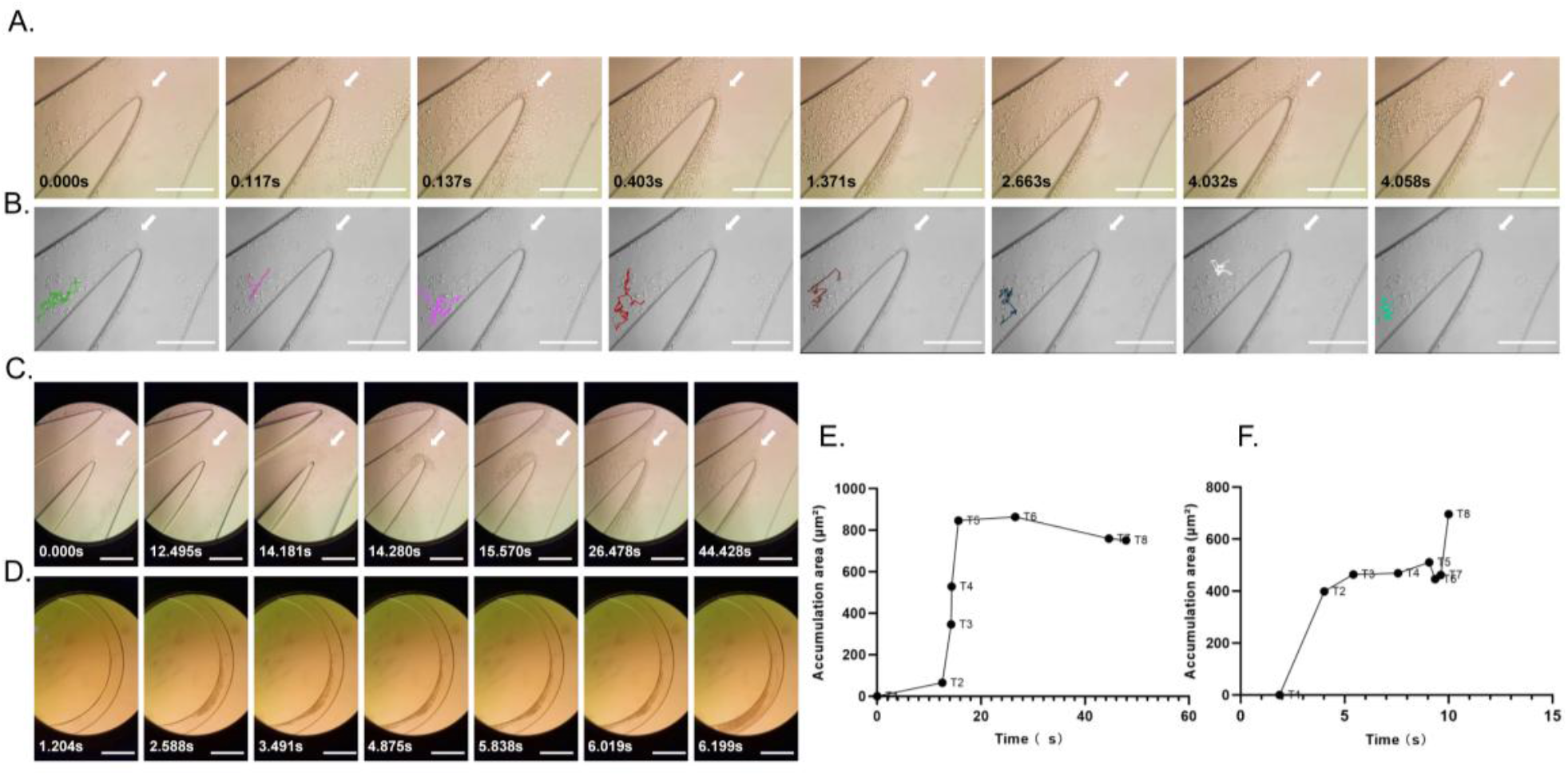
Flow-mediated spatiotemporal dynamics of tumor cell accumulation, adhesion, and mechanical remodeling at geometric confinements of a microfluidic chip. A. Representative micrographs illustrating cell accumulation and adhesion at the sharp corner, followed by flow-induced tensile stretching and elastic recoil of the viscoelastic, polymer-like network and cells. B. Migration trajectories of individual tracked cells. C. Time-lapse imaging showing the progressive accumulation and adhesion of tumor cells at the sharp corner of the microfluidic chip, with a time-dependent increase in the number of accumulated cells. D. Time-lapse imaging showing the progressive accumulation and adhesion of tumor cells along the inner wall of the curved channel in the microfluidic chip, with a time-dependent increase in the number of accumulated cells. E. Quantification of the accumulation area over time for the sharp-corner region shown in C. F. Quantification of the accumulation area over time for the curved-channel region shown in D. Data in E and F are from one representative experiment. The same trend was reproduced in three independent experiments (not shown). The accumulation area is reported in μm^2^.

**Figure 4.**
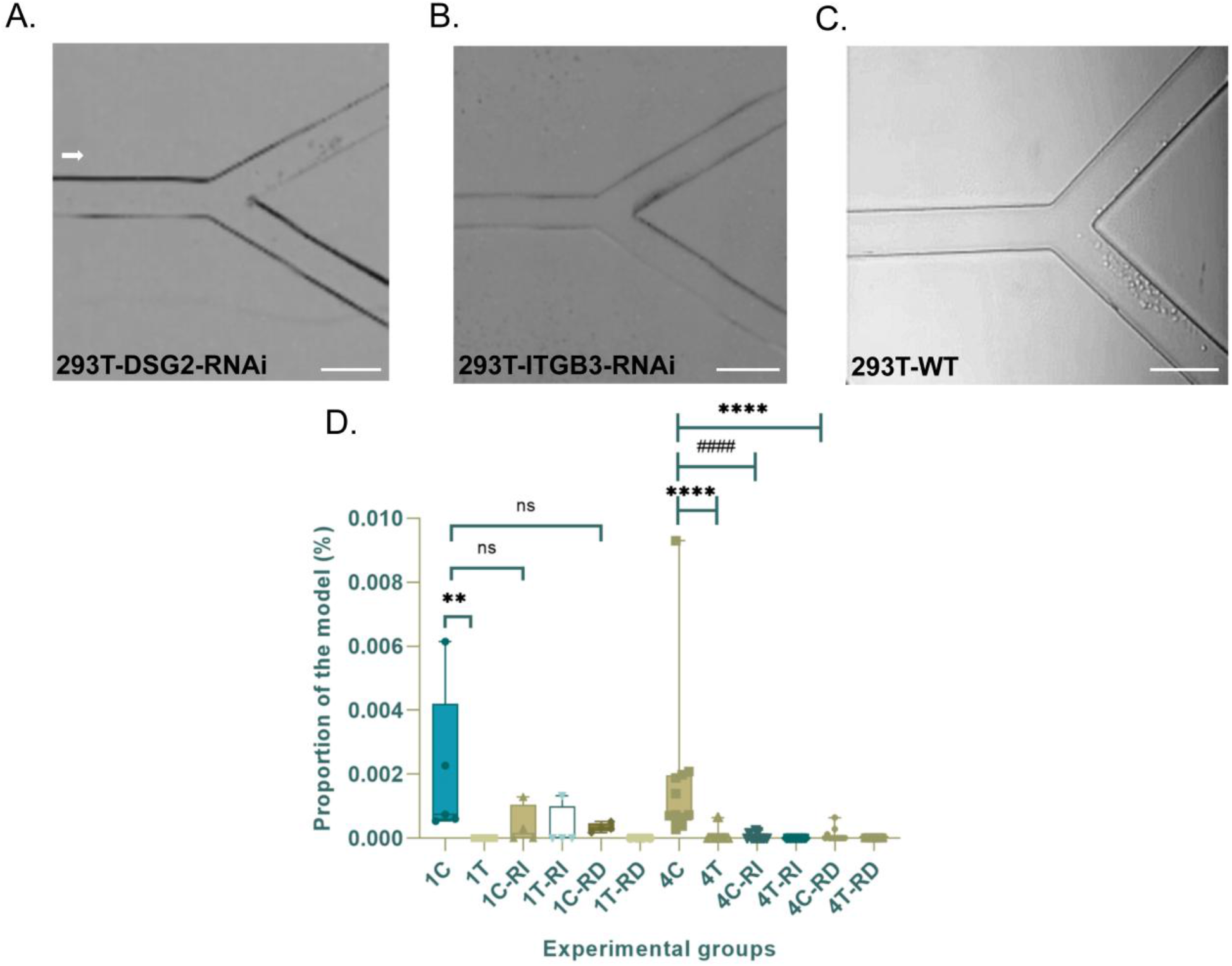
Mechanistic dissection of tumor-like cell accumulation, adhesion, and flow-induced multicellular aggregation at the sharp-corner region. A. Representative micrographs showing the accumulation and adhesion of DSG2-knockdown (DSG2-RNAi) tumor-like cells at the sharp corner. B. Representative micrographs showing the accumulation and adhesion of ITGB3-knockdown (ITGB3-RNAi) tumor-like cells at the sharp corner. C. Representative micrographs showing the accumulation and adhesion of wild-type (WT) tumor-like cells at the sharp corner. D. Quantitative analysis of tumor-like cell accumulation and adhesion in A–C. Data in D are presented as mean ± s.d. from n = 3 independent experiments; *P < 0.05, **P < 0.01, ***P < 0.001, ****P < 0.0001 (Kruskal−Wallis test followed by Dunn’s multiple comparisons test).

**Figure 5.**
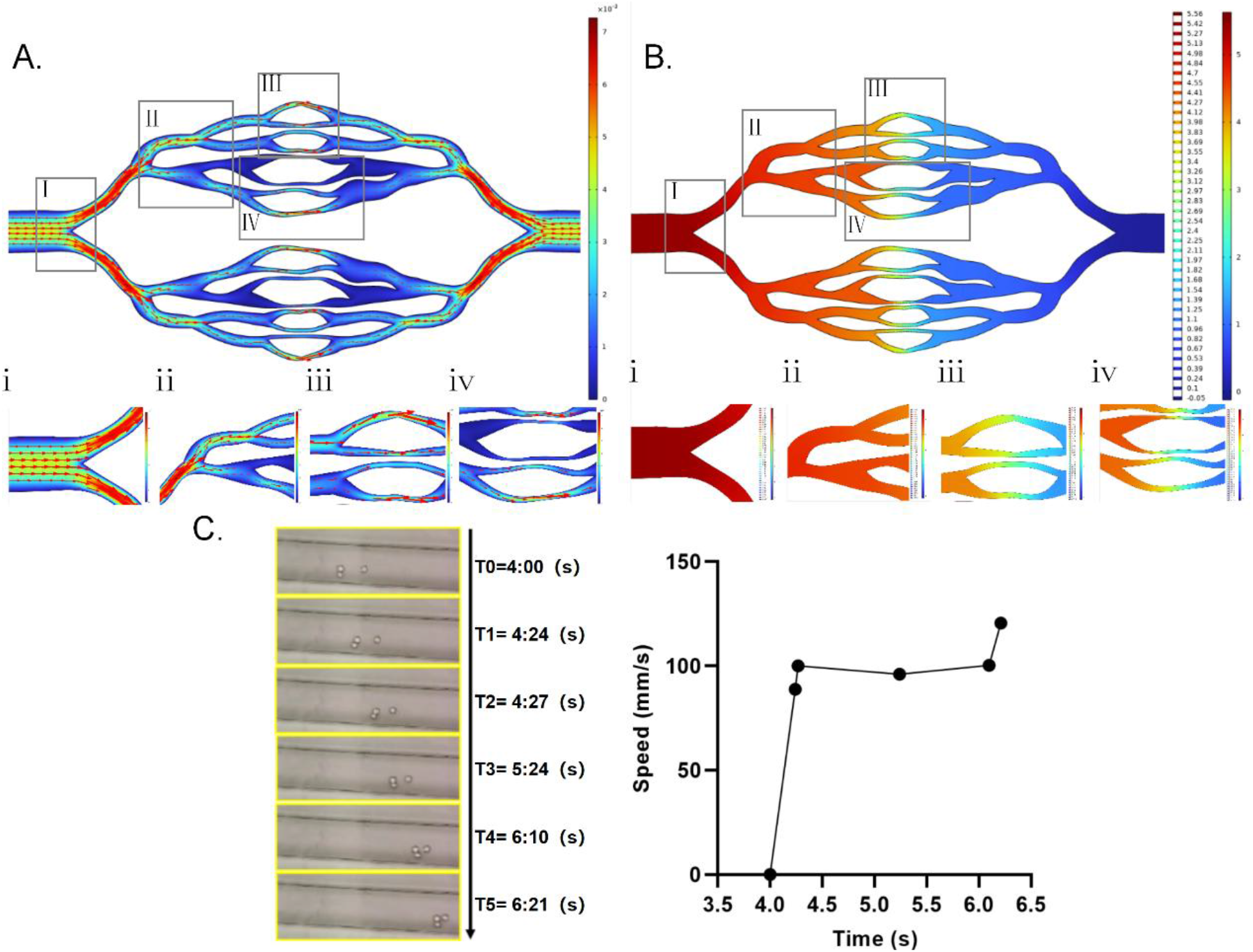
Computational fluid dynamics (CFD) simulations of the four-level vascular network performed with COMSOL Multiphysics, together with experimental imaging of flow-driven cell aggregation. A. Velocity magnitude distribution visualized as a heatmap, with arrows indicating flow direction and arrow size scaled to velocity magnitude. B. Pressure distribution. Regions of interest in A and B are labelled with uppercase Roman numerals (I–IV in A, I–IV in B), and the corresponding magnified views below each panel are labelled with the matching lowercase numerals (i–iv and i– iv, respectively). C. Left, representative time-lapse images showing progressive, flow-driven aggregation of tumor cells into multicellular clusters; right, quantification of tumor cell speed over time. Each data point represents a single measurement at the indicated time point. Similar trends were observed in independent experiments.

### Experimental

#### Cell culture

Two complementary cell backgrounds were selected to capture distinct aspects of circulating tumor cell (CTC) biology and to support live-cell imaging under flow. NCI-H69 human small-cell lung cancer cells served as a suspension model of CTCs. 293T human embryonic kidney cells further provided a robust cell-cell adhesion background for molecular perturbation. Both cell lines capture the essential characteristics of tumor cells – they circulate in the blood as suspended individual cells with weak adhesion to the substrate. All cells were incubated at 37 °C in a humidified atmosphere containing 5% CO_2_.

NCI-H69 cells were maintained in RPMI-1640 supplemented with 10% fetal bovine serum (FBS) and 2 mM L-glutamine. Because these cells grow in suspension, cultures were kept at 2–10 × 10^5^ cells/mL and passaged every 2–3 days by dilution or centrifugation–resuspension. Cell density and viability were monitored before each experiment to ensure consistent input for microfluidic perfusion.

293T cells were cultured in Dulbecco’s Modified Eagle Medium (DMEM) high glucose supplemented with 10% FBS, 2 mM L-glutamine, 1 mM sodium pyruvate, and non-essential amino acids. Cells were passaged at 70–80% confluence using 0.25% trypsin– EDTA and replated at 5 × 10^4^ cells/cm^2^ density.

Stable reporter and knockdown derivatives were propagated in the same base media and handled identically to their parental cells. Before microfluidic experiments, cells were harvested at log phase, washed, and resuspended in serum-free DMEM at 2 × 10^6^ cells/mL.

All cultures were routinely monitored for morphology, growth kinetics, and reporter expression. Mycoplasma contamination was excluded by fluorescence staining. Cell identity was confirmed by vendor certificate. Experiments were performed within 10 passages after thawing.

#### Construction of monoclonal cell lines

Lentiviral particles were produced using the lentiviral packaging kit (Genechem, Shanghai, China; cat. no. LPK002) according to the manufacturer’s protocol. Transfer plasmids encoding H2B-mCherry (histone H2B), a LifeAct-GFP actin reporter, desmoglein-2 (DSG2) short hairpin RNA (shRNA), integrin β3 (ITGB3) shRNA, or a non-targeting control were co-transfected with packaging plasmids into 293T cells. Viral supernatants were harvested and concentrated using the Takara lentiviral concentration kit (cat. no. 631231), yielding high-titer preparations suitable for efficient transduction of both adherent and suspension cells.

Target cells were transduced with concentrated virus. For suspension-grown NCI-H69 cells, transduction was performed in the presence of polybrene with spinoculation to enhance virus–cell contact, whereas adherent 293T cells were transduced under standard culture conditions. After 24 h, the medium was replaced to remove residual virus and polybrene. Antibiotic selection was then initiated and maintained for 3 days to eliminate untransduced cells and enrich the transduced population.

Single clones were isolated by limiting dilution and expanded in RPMI-1640 (NCI-H69) or DMEM high glucose (293T). Reporter expression and subcellular localization were verified by fluorescence microscopy. For NCI-H69 cells, clones expressing H2B-mCherry with a clear nuclear signal or LifeAct-GFP with a well-defined actin filament pattern were selected. For DSG2 and ITGB3 knockdown in 293T cells, each shRNA vector co-expressed a fluorescent reporter, enabling fluorescence-positive clones to be identified as knockdown candidates; reporter expression thereby served as a surrogate indicator of successful shRNA delivery and expression.

#### Fabrication of in vitro vessel networks and microfluidic devices

The microfluidic device was designed to reproduce the hierarchical architecture and geometric irregularities of tumor-associated vasculature, rather than a simplified straight channel. A four-level vascular network was laid out in computer-aided design (CAD), comprising a main vessel that sequentially branched into secondary vessels and capillary-like constrictions. Within this hierarchy, defined protruding geometries— sharp corners, curved segments, and stenosis-like narrowings—were intentionally incorporated to create controlled perturbations in the local flow field. This design allows cell behavior to be examined across multiple geometric scales within a single chip and permits direct comparison between straight segments and protruding confinements.

Masters were fabricated on silicon wafers by SU-8 photolithography using a single mask. The channel height was uniform at 100 μm. The four-level vascular network comprised a main vessel of ≈1.2 mm that branched progressively into narrower secondary vessels and capillary-like channels, the narrowest of which were ≈50–90 μm. After development, the wafers were silanized to facilitate mold release. Polydimethylsiloxane (PDMS) prepolymer and curing agent were mixed at 10:1, degassed under vacuum, poured over the master, and cured at 65 °C for 1 h. The cured PDMS replicas were peeled from the mold, and inlet and outlet ports were punched. Each replica was then bonded to a clean glass coverslip after oxygen plasma treatment, yielding optically transparent devices suitable for high-resolution bright-field, fluorescence, and confocal imaging.

Because all four hierarchical levels and the protruding features were defined in a single photolithographic step, the geometry of the perturbations was reproduced at the same positions in every replica, and devices were interchangeable between experiments. Each preparation therefore allowed cell behavior to be examined across the different geometric scales under controlled perfusion within a single chip.

#### Live-cell imaging

Live-cell imaging was performed on a perfusion-integrated platform housed within an incubator to maintain physiological temperature and CO_2_. The device was assembled as follows: the patterned PDMS layer and a clean glass coverslip were both exposed to oxygen plasma in a plasma cleaner (Shanghai Peiyuan Instrument Equipment Co., Ltd., Shanghai, China; model no. PLUTO-T) at 120 W for 1 min, brought into conformal contact, and bonded by baking at 65 °C for 1 h, yielding an irreversibly sealed, optically transparent channel network. The assembled device was reused across experiments. Between experiments, residual cells were detached by trypsin digestion for 15 min, followed by three phosphate-buffered saline (PBS) rinses and sonication for 30 min, until no residual material was detectable by microscopic inspection. Devices were then disinfected by soaking in 75% (v/v) ethanol overnight, soaked in sterile water for 2 h, and air-dried. Inlet and outlet ports were connected to tygon tubing (inner diameter 510 μm) through blunt stainless-steel needles. Before each experiment, the channel network was primed with cell-free culture medium to displace residual air and ensure bubble-free filling; the cell suspension prepared as described above (2 × 10^6^ cells/mL) was then drawn into a syringe and perfused through the inlet using a syringe pump (Infusetek, Shanghai, China; model no. SPM/2*AU-I) connected to the tubing. Perfusion was maintained at a constant volumetric flow rate of 20 μL min^-1^ for the duration of each recording (up to 3 h), and the corresponding flow velocities and wall shear stresses were estimated by CFD (see “Simulation of the flow field”).

The imaging system operated in two complementary modes—dynamic and static— each compatible with both bright-field and fluorescence channels. In the dynamic mode, a Huawei P40 smartphone (Huawei, Shenzhen, China; product no. ANA-AN00) coupled to a Yongxin inverted microscope (Ningbo Yongxin Optics Co., Ltd., Ningbo, China; model no. NIB910-FL) via a phone holder enabled continuous video recording of tumor cell accumulation, adhesion, migration, and multicellular aggregation under flow at 30 frames per second (fps; **Videos 1–4**). In the static mode, the motorized stage of the microscope, equipped with the Nomis Pro X software and its built-in stitching module, captured high-resolution images that were assembled into panoramic reconstructions of the entire chip. Bright-field imaging provided morphological context and large-field tracking, whereas fluorescence imaging reported the expression and localization of fluorescent reporter genes. At the experimental endpoint, the full-chip panorama provided a global spatial map of cell distribution across the four-level vascular network, allowing hotspots to be assigned to specific geometric features— bifurcations, sharp corners, curved segments, and stenosis-like constrictions—and enabling consistent comparison across devices. This dual-mode configuration— combining real-time dynamic recording with static panoramic mapping, and bright-field with fluorescence readouts—provided both temporal resolution and complete spatial coverage, overcoming the limited field of view of conventional live-cell microscopy.

#### Simulation of the flow field

To establish a mechanistic link between vascular geometry, local hemodynamics, and CTC homing, computational fluid dynamics (CFD) simulations were performed in COMSOL Multiphysics, resolving the hierarchical branching and in-plane protruding features—sharp corners, curved segments, and stenosis-like constrictions—identified experimentally as sites of preferential cell accumulation. The in-plane geometry was reconstructed from the CAD mask design and verified against volumetric imaging of the bonded device using a fluorescent tracer and Nomis Pro X software (Ningbo Yongxin Optics Co., Ltd., Ningbo, China). Because the channel depth was uniform at 100 μm and all geometric features were defined in the plane of the device, the flow field was approximated in two dimensions on the in-plane geometry using the Laminar Flow (spf) interface of COMSOL Multiphysics. In this formulation, the uniform depth was used to convert the volumetric flow rate into a flow rate per unit depth, so that the in-plane velocity field obtained from the two-dimensional solution approximates the depth-averaged velocity of the physical channel.

Flow was modeled as steady, laminar, incompressible, and Newtonian; the Reynolds number, evaluated from the hydraulic diameter of each channel level (D_h_ = 2wh/(w + h), where w and h are the channel width and depth, respectively), was ≈0.65 in the main vessel and below 1 throughout the device, confirming that inertial effects were negligible. The working fluid was cell culture medium (density 1000 kg m^-3^, dynamic viscosity 0.8 × 10^−3^ Pa s at 37 °C), matching the temperature at which live-cell experiments were performed. Boundary conditions comprised a fully developed inlet velocity profile corresponding to a rectangular channel with a flow rate per unit depth of 3.3 × 10^−6^ m^2^ s^-1^, corresponding to a total volumetric flow rate of 20 μL min^-1^ (mean velocity ≈2.8 mm s^−1^ in the ≈1.2 mm main vessel, with a peak in-plane velocity of ≈7.3 mm s^−1^ at the narrowest constrictions), a zero gauge pressure (atmospheric) outlet, and no-slip walls. Wall shear stress was estimated from the analytical relation for fully developed pressure-driven flow in a rectangular channel,τ = 6μQ/(wh^2^), giving 0.13 Pa (1.3 dyn cm^−2^) in the ≈1.2 mm main vessel. Because the flow divides among numerous parallel branches of differing widths, this analytical value serves as an order-of-magnitude estimate for the main vessel. To obtain the spatially resolved WSS in the physical 3D channel, a correction factor based on the local width-to-depth ratio was applied to the 2D CFD-derived wall shear rates. The steady-state two-dimensional incompressible Navier–Stokes equations were solved using a fully coupled damped Newton method with a direct (PARDISO) linear solver; convergence required scaled residuals below 1 × 10^−6^. The domain was discretized into approximately 4.4 × 10^5^ elements with local refinement near bifurcations, corners, and constrictions, and mesh independence was confirmed with the maximum velocity varying by less than 0.1% across successive refinements.

Post-processing focused on velocity magnitude and pressure distribution, visualized as heatmaps with arrows indicating flow direction and scaled to velocity magnitude (**Figure 5A**), and the corresponding pressure field (**Figure 5B**). Insets show magnified views of representative adhesion hotspots, enabling direct comparison between the simulated flow features and the experimentally observed accumulation sites. The simulated velocity and pressure fields were compared with the experimentally identified hotspots, and cell accumulation consistently coincided with the flow perturbations generated by the protruding geometry—reduced velocity, stagnation, and local pressure changes—indicating that these flow features underlie hotspot formation.

### Statistics and reproducibility

All measurements were obtained from n = 3 independent biological replicates across 10 independent devices. Data are presented as mean ± s.d. unless otherwise stated. Accumulation area (Figure 3E and F) was quantified from 316 cells per condition, and cell speed (Figure 5C, right) from 3 cells. Statistical significance was assessed using the Kruskal–Wallis test followed by Dunn’s multiple comparisons test (α = 0.05); *P < 0.05, **P < 0.01, ***P < 0.001, ****P < 0.0001. No data were excluded.

## Results and discussion

### Hotspots of circulating tumor cell homing exhibit protruding geometry

To determine whether the physical topography of the vasculature dictates where circulating tumor cells (CTCs) home, we perfused tumor cells through a multiscale microfluidic device that reproduces the hierarchical branching of a vascular network (**Figure 2A**). The chip design was adapted from the vascular-network platform reported by Pepona et al.^15^, and contained channels that transitioned from a main vessel to secondary branches and capillary-like constrictions, allowing cell behavior to be examined across multiple geometric scales. Throughout these experiments, flow was maintained under controlled conditions known to modulate CTC arrest and adhesion^7^.

Within a single-level vessel model, tumor cells were not distributed uniformly. Instead, they accumulated in discrete regions of the main channel, where adherent and aggregated cells were confined to defined locations rather than spread along the entire vessel length (**Figure 2B**). The insets highlight these representative hotspots, in which accumulation, adhesion, and aggregation coincided, suggesting that local geometric or hemodynamic conditions—rather than random cell–wall encounter—governed the initial stages of homing.

To resolve the structural basis of these hotspots, we examined branching points at higher resolution. Bright-field imaging showed that tumor cells accumulated and adhered preferentially at positions with protruded local geometry, such as bifurcations and corners (**Figure 2C, Video 1**). While bifurcations have been individually implicated in CTC arrest^9,16^, the preferential accumulation at corners represents a geometric feature not previously characterized in this context. Confocal imaging corroborated that adhesion and accumulation coincided at these same locations (**Figure 2D**), indicating that the observed hotspots reflected three-dimensional cell–wall interactions rather than projection artifacts. Notably, all hotspots were located at the upstream-facing surface rather than the downstream surface. This observation implies that geometry alone is not sufficient to cause CTC accumulation, indicating a coupled biophysical and biochemical underlying mechanism^8^.

Curved channel segments provided further evidence that protruding geometry shapes homing. Within a bend, tumor cells preferentially accumulated and adhered along the inner wall of the curvature (**Figure 2E, Video 2**), consistent with previous studies showing that tumor cell adhesion in curved microvessels occurs preferentially on the inner side of the curvature^17,18^. This behavior is consistent with the observation at bifurcations, because the inner wall of the bend likewise presents a protruding local contour.

Collectively, although bifurcations and curved segments have been separately associated with CTC arrest in prior work^9,16–18^, we propose that these sites share a common geometric signature—protruding or irregular contours that introduce local flow disturbances into an otherwise straight vessel. In this framework, corners represent an additional hotspot, not previously characterized in this context, belonging to the same geometric class. Together with the strictly upstream-facing localization of all hotspots observed here—which implies that geometry alone is insufficient and points to a coupled biophysical–biochemical mechanism^8^—this unifying geometric framework provides a coherent account of the observed hotspot distribution.

#### Viscoelastic, polymer-like network captures CTCs and facilitates their accumulation

Having established that geometric protrusions serve as homing hotspots, we next asked how tumor cells are retained at these sites. To address this, we examined the spatiotemporal dynamics of cell accumulation and the mechanical behavior of the associated actin cytoskeleton and cell-derived structures at a sharp corner and along the inner wall of a curved channel (**Figure 3**).

At the sharp corner, tumor cells accumulated and adhered progressively. In cells labeled with LifeAct-GFP, actin cytoskeleton-associated structures, together with the cells themselves, underwent flow-induced tensile stretching followed by elastic recoil (**Figure 3A, Video 1**). This stretch–recoil behavior is consistent with the mechanics of a viscoelastic, polymer-like material^19,20^, raising the possibility that such a network assembles at geometric confinements and responds mechanically to local flow. Hereafter, this actin cytoskeleton-associated protein/ECM assembly is referred to as the viscoelastic, polymer-like network. Tracking of individual cells revealed that their migration trajectories were constrained near the corner, consistent with capture by this network rather than free migration (**Figure 3B**).

Time-lapse imaging confirmed that accumulation at the sharp corner was progressive, with the number of accumulated cells increasing over time (**Figure 3C, Video 1**). A similar time-dependent increase was observed along the inner wall of the curved channel (**Figure 3D, Video 2**). In both geometries, cells were retained at the same locations where the cytoskeleton-associated structures—extending beyond the reach of single-cell contacts—were mechanically deformed. Although hemodynamic profiles are known to tune CTC arrest through a balance of flow and adhesion forces at vascular junctions^7,9,21^, our observations point to a distinct, network-mediated mechanism: these structures appear to capture CTCs and sustain their accumulation over distances far exceeding those expected from single-cell contact, a long-range capture mechanism that single-cell adhesion cannot explain and that is consistent with geometry-induced enhancement of cell–wall contact probability^8,21^. This long-range feature significantly enhances CTC capture efficiency and contact probability with the inner wall of blood vessels, thereby promoting homing and even metastasis^7,22^.

Collectively, these observations suggest that geometric confinements promote the assembly of a viscoelastic, polymer-like network, whose deformation—manifested in the stretch–recoil behavior of the associated actin cytoskeleton—coincides with the capture and progressive accumulation of circulating tumor cells. This mechanism provides a mechanical explanation for the hotspot formation observed in the hierarchically branched vascular network^7,21^.

#### Mechanistic analysis: molecular and mechanical coupling

Having established that geometric confinements capture CTCs through a viscoelastic, polymer-like network, we next dissected the molecular and mechanical basis of this process. To identify the molecular players involved, we knocked down two candidate adhesion-associated molecules by RNA interference (RNAi), DSG2^23^ and ITGB3 (the β3 subunit of αvβ3 integrins)^24^, and compared their accumulation and adhesion behavior at the sharp corner with that of wild-type (WT) cells (**Figure 4A–C, Video 3**). Quantitative analysis revealed that both DSG2-RNAi and ITGB3-RNAi cells exhibited reduced accumulation and adhesion relative to WT cells (**Figure 4D**), indicating that DSG2 and ITGB3 contribute to cell retention at geometric confinements. This is consistent with previous work showing that loss of DSG2 reduces tumor cell cluster adhesion during vascular perfusion^25^ and that integrin activation supports CTC arrest under flow^24^.

To determine whether the experimentally observed hotspots correspond to regions of favorable hemodynamics, we performed computational fluid dynamics (CFD) simulations using COMSOL Multiphysics on a custom-designed four-level branched network whose in-plane geometry was reconstructed from the CAD layout of the device (**Figure 5A, B**). The design was conceptually informed by classical microvascular hemodynamics^26^, and key geometric parameters and boundary conditions are provided in Supplementary Information. The simulated velocity and pressure fields revealed that the sharp-corner and curved regions experience pronounced flow perturbations, consistent with the experimental hotspots of cell accumulation and adhesion. While hemodynamic profiles are known to tune CTC arrest and adhesion^7^, our simulations reveal that geometric confinement locally reshapes these profiles in a manner that spatially coincides with the experimental hotspots.

We then examined the dynamics of cell–cell association under flow. Time-lapse imaging showed that tumor cells progressively aggregated into multicellular clusters in a straight channel (**Figure 5C, left, Video 4**), consistent with extracellular matrix (ECM)-mediated CTC clustering that enables survival under flow^27^; such clusters are recognized as potent metastatic precursors^28^. Whereas ECM-mediated clustering in the circulation is not spatially restricted^27^, the accumulation of the resulting clusters observed here is spatially confined to geometric hotspots and is coupled to the viscoelastic network. Quantification of cell speed over time in the same segment confirmed that cells were transported by the local flow (**Figure 5C, right**). These observations suggest that the accumulation of multicellular clusters at geometric hotspots involves ECM-mediated cell–cell contacts acting in concert with local flow, beyond single-cell adhesion^27^.

Collectively, these results indicate that CTC accumulation at geometric confinements is governed by a coupled molecular and mechanical mechanism: DSG2 and ITGB3 mediate cell–cell and cell–matrix adhesion^23,24^, while flow-induced velocity and pressure fields drive cells toward these regions^7,26^ and ECM-based cell–cell contacts stabilize the resulting clusters^27^.

This study identifies vascular geometry as a decisive determinant of where CTCs arrest and accumulate. Using a multiscale microfluidic device that reproduces hierarchical vascular branching, we show that CTC homing concentrates at protruding geometric confinements—bifurcations, sharp corners, and curved segments—rather than being randomly distributed. Whereas previous studies examined individual geometric elements in simplified vessel segments^9,17^, here the emergence of these features within a hierarchically branching network, and their relative arrangement, is associated with arrest. Notably, the viscoelastic protein/ECM network that assembles at these confinements sustains CTC accumulation over distances far exceeding the reach of single-cell receptor–ligand contacts. This long-range capture substantially increases the probability of cell–wall contact and retention, providing a geometrically amplified mechanism for CTC homing that is distinct from conventional single-cell adhesion models^7,8,21^.

Mechanistically, this capture arises from a biochemical–mechanical coupling. Geometric confinements promote assembly of a viscoelastic, polymer-like protein/ECM network whose mechanical remodeling—manifested as stretching and elastic recoil—actively retains cells. Molecular dissection supports this coupling: knockdown of DSG2 or ITGB3 reduced accumulation and adhesion, implicating DSG2 in cell–cell adhesion^23,29,30^ and ITGB3 in cell–substrate interaction^24,31,32^. These findings suggest potential intervention strategies targeting the long-range polymer network and its cytoskeletal remodeling (e.g., actin dynamics), cell–cell adhesion (DSG2), and cell–substrate interaction (ITGB3) to reduce CTC accumulation^31,33^.

CFD simulations indicated that experimental hotspots coincide with pronounced velocity and pressure perturbations^34^, showing that geometric confinement locally reshapes flow in a manner spatially coinciding with cell accumulation. This integration of multiscale microfluidics, live-cell imaging, molecular perturbation, and CFD provides a template for studying other blood-vessel processes such as immune cell trafficking^35^ and thrombus formation^36^. Limitations include the lack of endothelialization, so tumor cell–endothelium interactions require further study^4^, and the unquantified composition and rheology of the network^19,20^. Future work should incorporate endothelial cells and co-culture stromal cells or ECM to better mimic the metastatic niche^37^.

## Conclusions

Vascular geometry, rather than flow alone, determines where circulating tumor cells arrest and accumulate. In a multiscale microfluidic device reproducing the hierarchical branching of a vascular network, protruding and positively curved contours introduced discrete locations at which tumor cells were captured, and these sites coincided with the controlled perturbations in the flow field. At these hotspots, a viscoelastic, polymer-like protein/ECM network underwent flow-induced stretching and elastic recoil and progressively retained cells, while knockdown of DSG2 or ITGB3 reduced accumulation and adhesion—revealing that local mechanical remodeling and molecular adhesion act in concert. Computational fluid dynamics simulations independently indicated that the experimental hotspots coincided with pronounced velocity and pressure perturbations. Together, these results identify local geometry as a critical and previously underappreciated determinant of CTC homing, and establish a platform for dissecting the physical mechanisms of metastatic seeding. Extending the system to endothelialized and co-cultured channels represents the natural next step.

## Acknowledgments

Bo Li acknowledges the support from taxpayers of China through the National Natural Science Foundation of China (Distinguished Young Scholars Funding, Overseas; Young Scientists Fund – Category C, Grant No. 32500692) and Wuhan University (Talents Startup Funding). DeepSeek V4 was used for text polishing and grammar checking under the supervision of the authors.

## Author contributions

B. L. conceptualized the project and designed the research. Y. Q. L. performed experiments and analyzed the data. All authors discussed the results and wrote the paper.

## Data availability

The data and analysis code that support the findings of this study are available upon request to the corresponding author.

## Video captions

**Video 1 Flow-induced stretching and elastic recoil of the actin cytoskeleton-associated polymer-like network**. Left, bright-field imaging; right, fluorescence imaging of the same field of view. Cells were labeled with LifeAct-GFP and imaged in the green channel (488 nm). The actin cytoskeleton-associated structures, together with the cells themselves, undergo flow-induced tensile stretching followed by elastic recoil, consistent with the mechanics of a viscoelastic, polymer-like network. Recorded with a Huawei P40 smartphone camera at 30 frames per second (fps) and shown at 5× speed.

**Video 2 Accumulation of circulating tumor cells along a curved channel segment under flow**. Left, bright-field imaging; right, fluorescence imaging of the same field of view. Cells were labeled with H2B-mCherry and imaged in the red channel (561 nm). The recording shows preferential accumulation and adhesion of CTCs along the inner wall of a curved channel segment under continuous perfusion. Recorded with a Huawei P40 smartphone camera at 30 fps and shown at real time.

**Video 3 Accumulation of tumor-like cells under flow in DSG2-RNAi and ITGB3-RNAi cells**. Bright-field recording of CTC accumulation and adhesion within the microfluidic channel. DSG2-RNAi cells are shown on the left and ITGB3-RNAi on the right. Both conditions were perfused under the same controlled flow and recorded in parallel. Recorded with a Huawei P40 smartphone camera at 30 fps and shown at real time. Note the reduced accumulation and adhesion in both RNAi conditions relative to wild-type cells (Figure 4D).

**Video 4 Progressive aggregation of circulating tumor cells into multicellular clusters under flow**. Recording of CTCs progressively aggregating into multicellular clusters under continuous perfusion in a straight channel segment. Tracking arrows are overlaid to follow the cells as they aggregate. Recorded with a Huawei P40 smartphone camera at 30 fps and shown at 0.3× speed.

